# Building a 3D Integrated Cell

**DOI:** 10.1101/238378

**Authors:** Gregory R. Johnson, Rory M. Donovan-Maiye, Mary M. Maleckar

## Abstract

We present a conditional generative model for learning variation in cell and nuclear morphology and predicting the location of subcellular structures from 3D microscopy images. The model generalizes well to a wide array of structures and allows for a probabilistic interpretation of cell and nuclear morphology and structure localization from fluorescence images. We demonstrate the effectiveness of the approach by producing and evaluating photo-realistic 3D cell images using the generative model, and show that the conditional nature of the model provides the ability to predict the localization of unobserved structures, given cell and nuclear morphology. We additionally explore the model’s utility in a number of applications, including cellular integration from multiple experiments and exploration of variation in structure localization. Finally, we discuss the model in the context of foundational and contemporary work and suggest forthcoming extensions.

## 1. Introduction

A central conjecture throughout cell biology is that structure determines function. Thus motivated, location proteomics (Murphy, 2005) aims to determine cell state – i.e. subcellular organization – by elucidating the localization of *all* structures and how they change through the cell cycle, and in response to perturbations in environment or mutation, for example. However, determining global cellular organization is challenging, not in small part by the multitude of different molecules, complexes and organelles that comprise living cells and determine their behaviors (Kim et al., 2014). While advances in microscopy, particularly live cell fluorescence imaging, have permitted enormous insight and rich datasets with which to explore subcellular organization, the experimental state-of-the-art for live cell imaging is currently limited to the simultaneous visualization of only a limited number of tagged (2-6 tagged) molecules. Statistical modeling approaches can address this limitation by integrating subcellular structure data from diverse imaging experiments.

Image feature-based methods have previously been employed to describe and model cell organization (Boland & Murphy, 2001; Carpenter et al., 2006; Rajaram et al., 2012). While useful for discriminative tasks, these approaches do not explicitly model the relationships among subcellular components, limiting their use to integration of all of these structures, i.e., creating an integrated cell model.

Generative models are useful in this context.They capture variation in a population and encode it as a probability distribution, accounting for the relationships among structures. Fundamental work has previously demonstrated the utility of expressing subcellular structure patterns as a generative model, which can then be used as a building block for models of cell behavior, i.e. (Murphy, 2005; Donovan et al., 2016).

Ongoing efforts to construct generative models of cell organization are primarily associated with the CellOrganizer project (Zhao & Murphy, 2007; Peng & Murphy, 2011). CellOrganizer implements a “cytometric” approach to modeling that considers the number of objects, lengths, sizes, etc. from segmented images and/or inverse procedural modeling, which can be particularly useful for both analyzing image content and approaching integrated cell organization. These methods support parametric modeling of many subcellular structure types and, as such, generalize well when low amounts of appropriate imaging data are available. However, these models may depend on preprocessing methods, such as segmentation, or other object identification tasks for which a ground truth is not available. Additionally, there may exist subcellular structures for which a parametric model does not exist or may not be appropriate e.g., structures that vary widely in localization (diffuse proteins), or that reorganize dramatically during e.g. mitosis or during a stimulated state (such as microtubules).

Thus, the presence of key structures for which current methods are not well suited motivates the need for a new approach that generalizes well to a wide range of structure localization.

Recent advances in generative adversarial networks (GANs) (Goodfellow et al., 2014) present one possible resolution to this dilemma. GANs have the ability to learn distributions over images, generate photo-realistic exemplars, and learn sophisticated conditional relationships; see e.g. Generative Adversarial Networks (Goodfellow et al., 2014), Varational Autoencoders/GAN (Larsen et al., 2015), Adversarial Autoencoders (Makhzani et al., 2015).

Leveraging these advances, we present a non-parametric model of 3D cell shape and nuclear shape and location, and relate it to the variation of other subcellular components. The model is trained on data sets of approximately 160–4400 fluorescence microscopy images per labeled structure; it accounts for the spatial relationships among these components, their fluorescent intensities, and generalizes well to a variety of localization patterns. Using these relationships, the model allows us to predict the outcome of theoretical experiments, as well as encode complex image distributions into a low dimensional probabilistic representation. This latent space serves as a compact coordinate system to explore variation.

Here, we present the integrated cell model, a discussion of the training and conditional modeling, and initial results demonstrating its utility. We then briefly discuss the results in context, current limitations of the work and future extensions.

## 2. Model Description

Our generative model serves several distinct but complementary purposes. At its core, it is a probabilistic model of cell and nuclear shape (specifically, of cell shape and nuclear shape and *location*) conjoined with a probability distribution for the localization of a given fluorescent protein (for these experiments, proteins were chosen to outline major cellular structures) conditional on cell and to classify localization patterns from images where the protein is unknown, and to predict the localization of unobserved structures *de novo*.

The main components of the model (Figure 1) are two modified autoencoders; one which encodes the variation in cell and nuclear shape (reference structure model), and another which learns the relationship between subcellular structures dependent on this encoding (conditional model). Each autoencoder is regularized by the use of adversarial networks on the latent space (to enforce the distribution of latent space representations matches a known distribution), and on the output (to enforce that *de novo* generated images match the data distribution).

**Figure 1:**
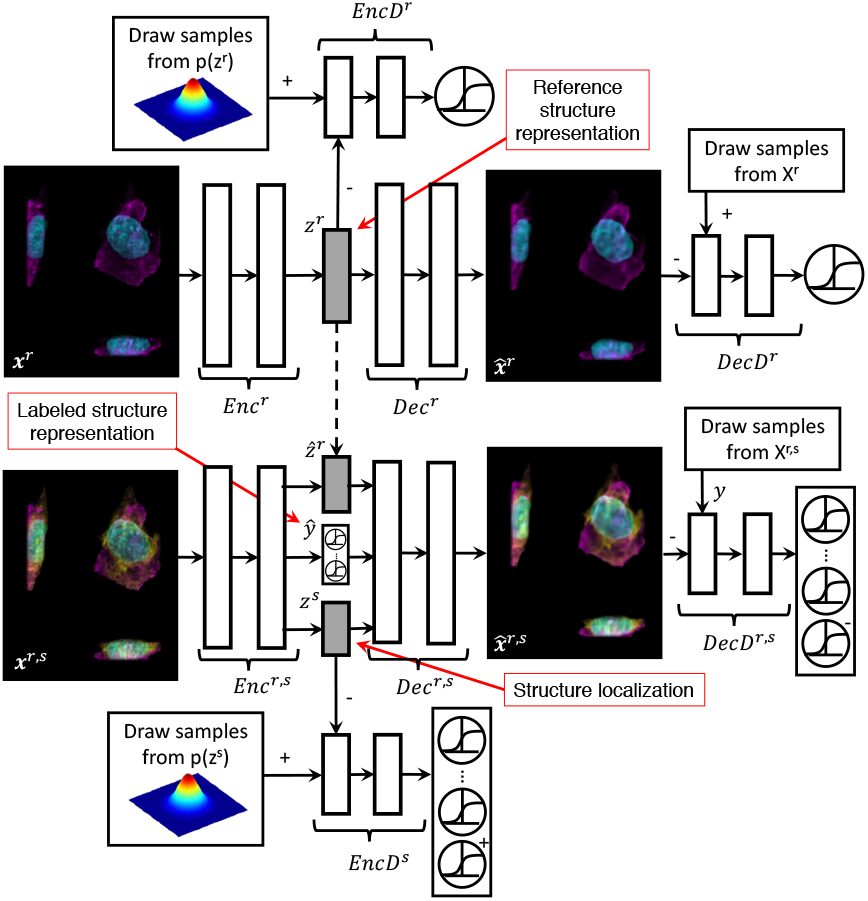
Conditional model of subcellular localization. The top half of the diagram outlines the reference structure model; the bottom half shows the conditional model. The parallel white boxes indicate a nonlinear function. The model is a probabilistic model of cell and nuclear shape (specifically, of cell shape and nuclear shape and *location*, the reference structure model) wedded to a probability distribution of structure localization (e.g. the localization of a certain protein) conditional on cell and nuclear shape, the conditional model. This model can be used both as a classifier for images of localization pattern where the protein is unknown, and as a tool for prediction of the localization of unobserved structures *de novo*. The main components are two autoencoders: one encoding the variation in cell and nuclear shape, and another which learns the relationship between subcellular structures dependent on this encoding. See Notation and Model description for details. Figure adapted from (Makhzani et al., 2015)

### Notation

The images input and output by the model are multi-channel (Figure 2). Each image *x* consists of both reference channels *r* and a structure channel *s*. Here, the cell and nuclear channels together serve as reference channels, and the structure channel varies, taking on one of the following structure types: α-actinin (actin bundles), α-tubulin (microtubules), β-actin (actin filaments), desmoplakin (desmosomes), fibrillarin (nucleolus), sialyltransferase (golgi), lamin B1 (nuclear membrane), myosin IIB (actomyosin bundles), Sec61β(endoplasmic reticulum), TOM20 (mitochondria), and ZO1 (tight junctions). We denote which content is being used by superscripts; ***x**^r,s^* indicates all channels are being used, whereas ***x**^s^* indicates only the structure channel is being used, and ***x**^r^* only the reference channels. We use *y* to denotes an index-valued categorical variable indicating which structure type is labeled in ***x**^s^*. For example, *y* = 1 might correspond to the α-actinin channel being active, *y* = 2 to the α-tubulin channel, etc. While *y* is a scalar integer, we also use *y*, a one-hot vector representation of *y*, with a one in the *y*th element of *y* and zeros elsewhere.

**Figure 2:**
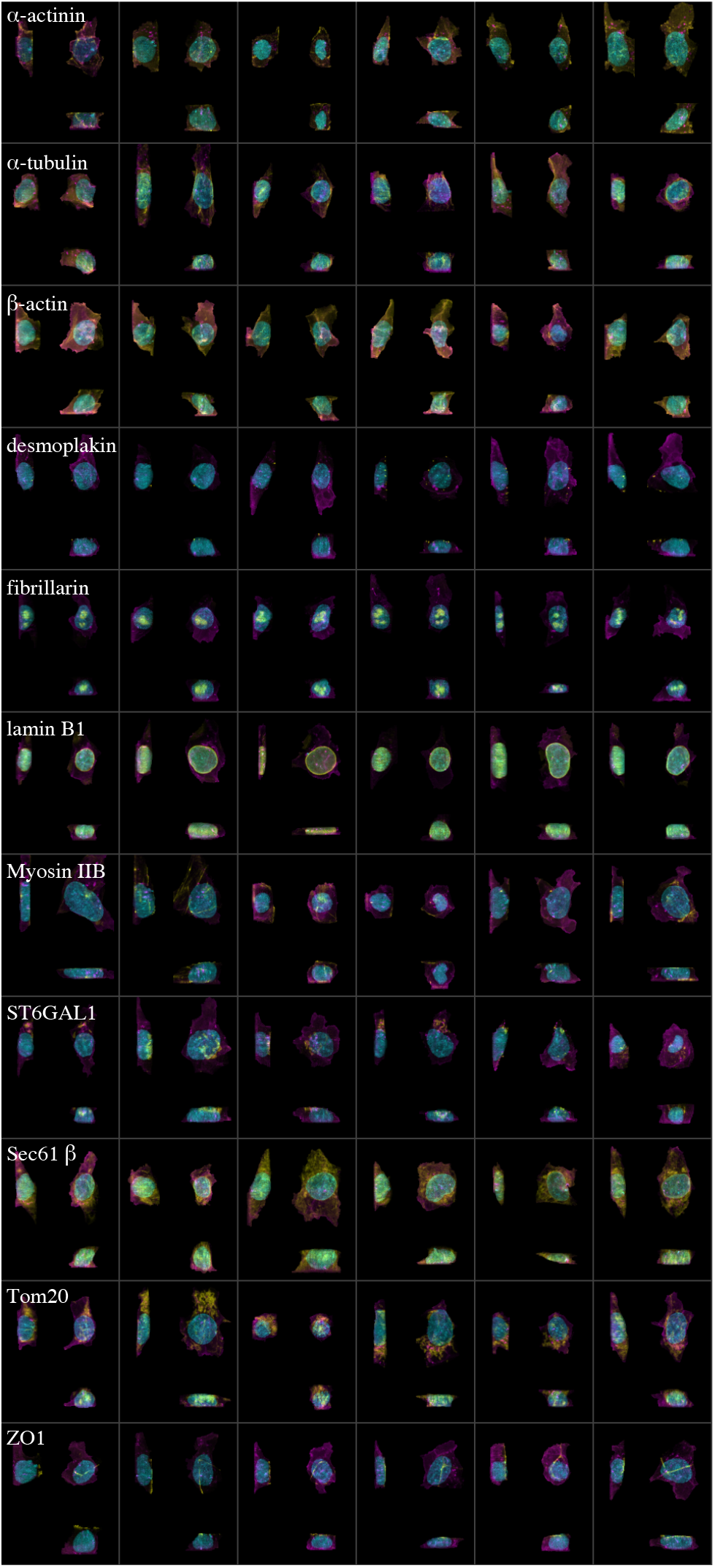
Example images for each of the 10 labeled structures of interest. Rows correspond to observed microscopy images, used as inputs to the model, for six arbitrary cells, each with a particular fluorescently labeled structure as named, shown in yellow. The reference structures, the cell membrane and nucleus (DNA), are shown in magenta and cyan, respectively. Images have been cropped for visualization purposes. See Figure S5a for isolated observed structure channel only.

### 2.1 Model of cell and nuclear variation

We model cell and nuclear shape using an autoencoder to construct a latent-space representation of these reference channels. The model (Figure 1, upper half) maps images of reference channels to a multivariate normal distribution of moderate dimension – here we use a 128 dimensional distribution. The choice of a normal distribution as the prior for the latent space is, in many respects, one of convenience, and of no large consequence to model behavior. The nonlinear mappings learned by the encoder and decoder are coupled to both the shape and dimensionality of the latent space distribution; the mapping and the distribution only function in tandem – see e.g. Figure 4 in (Makhzani et al., 2015).

The primary architecture of the model is that of an autoencoder, which itself consists of two networks: an encoder Enc*^r^* that maps an image ***x*** to a latent space representation *z* via a learned deterministic function *q*(*z^r^*|***x**^r^*), and a decoder Dec*^r^* to reconstruct samples from the latent space representation using a similarly learned function 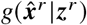.

We use the following notation for these mappings:

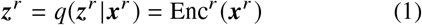

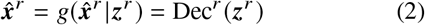

where an input image ***x*** is distinguished from a reconstructed image 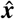.

#### 2.1.1. Encoder and Decoder

The autoencoder minimizes the pixel-wise binary crossentropy loss of the input and reconstructed input using binary cross entropy,

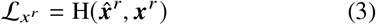

where

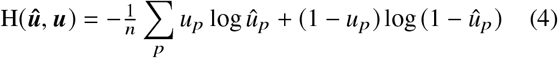

and the sum is over all the voxels *p* in all the channels in the images ***u***. We use this function for all images regardless of content (i.e. we use it for ***x**^r^* and ***x**^r,s^*)

#### 2.1.2. Encoding Discriminator

In addition to minimizing the above loss function, the autoencoder’s latent space – the output of Enc*^r^* – is regularized by the use of an encoding discriminator EncD*^r^*. This discriminator EncD*^r^* attempts to distinguish between latent space embeddings that are mapped from the input data, and latent space embeddings that are generatively drawn from the desired prior latent space distribution (which is a 128 dimensional multivariate normal in this case). In attempting to “fool” the discriminator, the autoencoder is forced to learn a latent space distribution *q*(*z^r^*) that is similar in distribution to the prior distribution *p*(*z^r^*) (Makhzani et al., 2015).

The encoding discriminator EncD*^r^* is trained on samples from both the embedding space *z* ~ *q*(*z^r^*) and from the desired prior 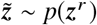. We refer to *z* as observed samples, and 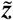 as generated samples, and use the subscripts “obs” and “gen” to indicate these labels. Trained on these samples, EncD*^r^* outputs a continuous estimate of the source distribution, 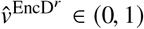.

The objective function for the encoding discriminator is thus to minimize the binary-cross entropy between the true labels *v* and the estimated labels 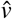 for generated and observed images:

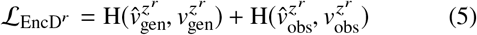

#### 2.1.3. Decoding Discriminator

The final component of the autoencoder for cell and nuclear shape is an additional adversarial network DecD *r*, the decoding discriminator, which operates on the output of the decoder to ensure that the decoded images are representative of the observed data distribution, similar to that of (Larsen et al., 2015). We train DecD *r* on images from the data distribution, 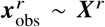, which we refer to as observed images, and on decoded draws from the latent space, 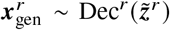, which we refer to as generated images. The loss function for the decoding discriminator is then:

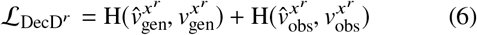

### 2.2 Conditional model of structure localization

Given a trained model of cell and nuclear shape variation from the above network component, we then train a conditional model of structure localization upon the learned cell and nuclear shape model. This model (Figure 1, lower) consists of several parts, analogous to those outlined in the previous section: the core is a tandem encoder Enc*^r,s^* and decoder Dec*^r,s^* that encode and decode images to and from a low dimensional latent space; in addition, a discriminative decoder EncD*^s^* regularizes the latent space, and a discriminative decoder DecD*^r,s^* ensures that the decoded images are similar to the input distribution.

#### 2.2.1. Conditional Encoder

The encoder Enc*^r,s^* is given images containing both the reference structures and a single additional structure of tagged protein localization, ***x**^r,s^*, and produces three outputs:

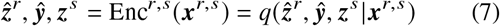

Here 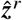 is the reconstructed cell and nuclear shape latent-space representation learned in Section 2.1, ***ŷ*** is an estimate of which structure channel was learned, and *z^s^* is a latent variable that encodes all remaining variation in image content not due to cell/nuclear shape and structure channel. Therefore *z^s^* is learned dependent on the latent space embeddings of the reference structure, *z^r^*.

The loss function for the reconstruction of the latent space embedding of the cell and nuclear shape is the mean squared error between the embedding *z^r^* learned from the cell and nuclear shape autoencoder and the estimate 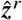 of that embedding produced by the conditional portion of the model:

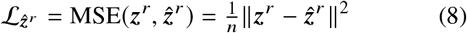

The output ***ŷ*** in equation7 is a probability distribution over structure channels, giving an estimate of the class label for the structure. In our notation, *y* is an integer value representing the true structure channel, and takes an integer value 1…*K*, while *y* is the one-hot encoding of that label, a vector of length *K* equal to 1 at the *y*th position and 0 otherwise. Similarly, *ŷ* is a vector of length *K* whose *k*th element represents the probability of assigning the label *y* = *k*.

We use the softmax function to assign these probabilities. In general, the softmax function is given by

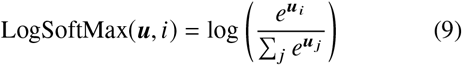

The loss function for ***ŷ*** is then

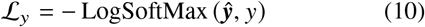

The final output of the conditional encoder *z^s^* can be interpreted as a variable that encodes the residual variation in the localization of the labeled structure after conditioning on cell and nuclear shape.

#### 2.2.2. Encoding Discriminator

As introduced above, the latent variable *z^s^* is regularized by an adversary EncD*_s_* that enforces the distribution of this latent variable to be similar to a chosen prior *p*(*z^s^*). The output of EncD*_s_* is a vector ***ŷ***^EncD^*s*^^ that has |***y***| + 1 = *K* + 1 output labels, which take a value in [1,…, *K*, gen]. That is, ***ŷ***^EncD^*s*^^ has one slot for observed embeddings of each particular labeled structure channel, and one additional slot for samples from our reference distribution. The loss function for the adversary is therefore:

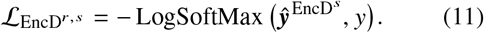

The construction of the loss function is intended to remove structure-specific information from *z^s^*, forcing that information into *ŷ*.

#### 2.2.3. Conditional Decoder

The conditional decoder Dec*^r,s^* outputs the image reconstruction given the latent space embedding *z^r^*, the class estimator ***ŷ***, and the structure channel variation *z^s^*:

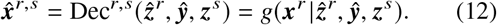

The loss function for image reconstruction takes the same form as equation3 (the binary cross entropy between the input and reconstructed image):

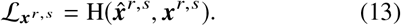

#### 2.2.4. Decoding Discriminator

As in the cell and nuclear shape model, attached to the decoder Dec*^r,s^* is an adversary DecD*^r,s^* intended to enforce that the reconstructed images are similar in distribution to the input images. Similar to Section 2.2.2, the output of this discriminator is a vector ***ŷ***^DecD*^r,s^*^ that has |***y***| + 1 = *K* + 1 output labels, which take a value in [1,…, *K*, gen]. As above, the loss function is:

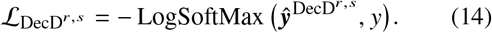

### 2.3 Training procedure

The training procedure occurs in two phases. We first train the model of cell and nuclear shape variation, components Enc*^r^*, Dec*^r^*, EncD*^r^*, DecD*^r^* to convergence (Algorithm 1). We then train the conditional model, components Enc*^r,s^*, Dec*^r,s^*, EncD*^s^*, DecD*^r,s^* (Algorithm 2).

In training the model, we adopt three strategies from (Larsen et al., 2015): we limit error signals to relevant networks by propagating the gradient update from any DecD through only Dec, we train the image adversary, DecD, with both generated and reconstructed images, and we weight the gradient update from the discriminators with the scalars *γ*_Enc_ and *γ*Dec. The parameters of the model are therefore updated as follows:

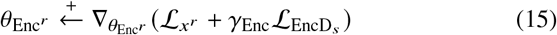

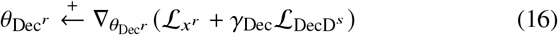

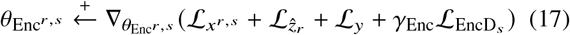

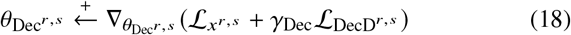

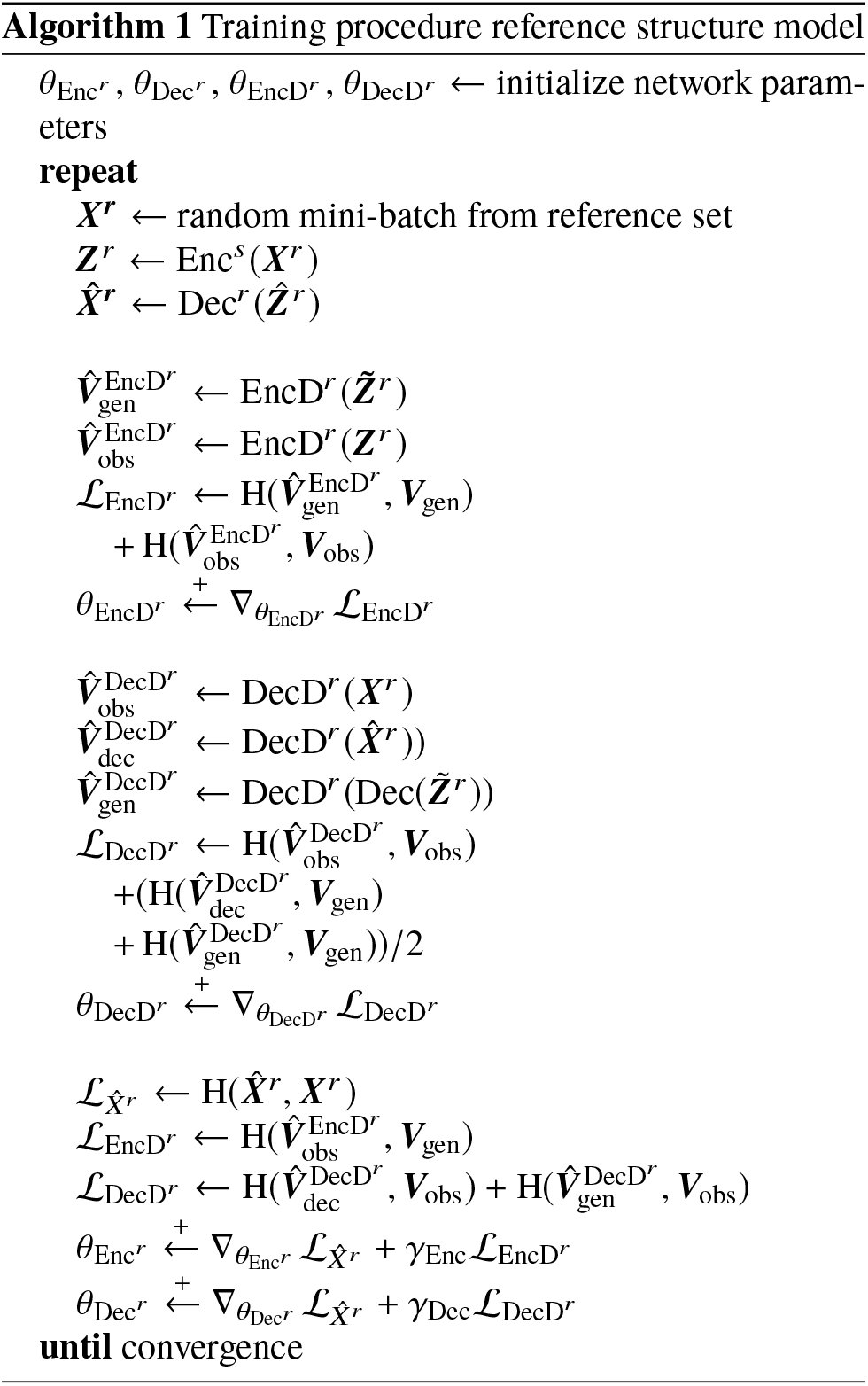

### 2.4 Integrative Modelling

Beyond encoding and decoding images, we leverage the conditional model of structure localization given cell and nuclear shape (Section 2.1) as a tool to predict the localization of multiple unobserved structures, *p*(*x^s^*|*x^r^*, *y*), in a known cell and nuclear representation. In particular, we predict using the most probable structure localization given the cell and nuclear channels. The procedure for predicting this localization is outlined in Algorithm 3.

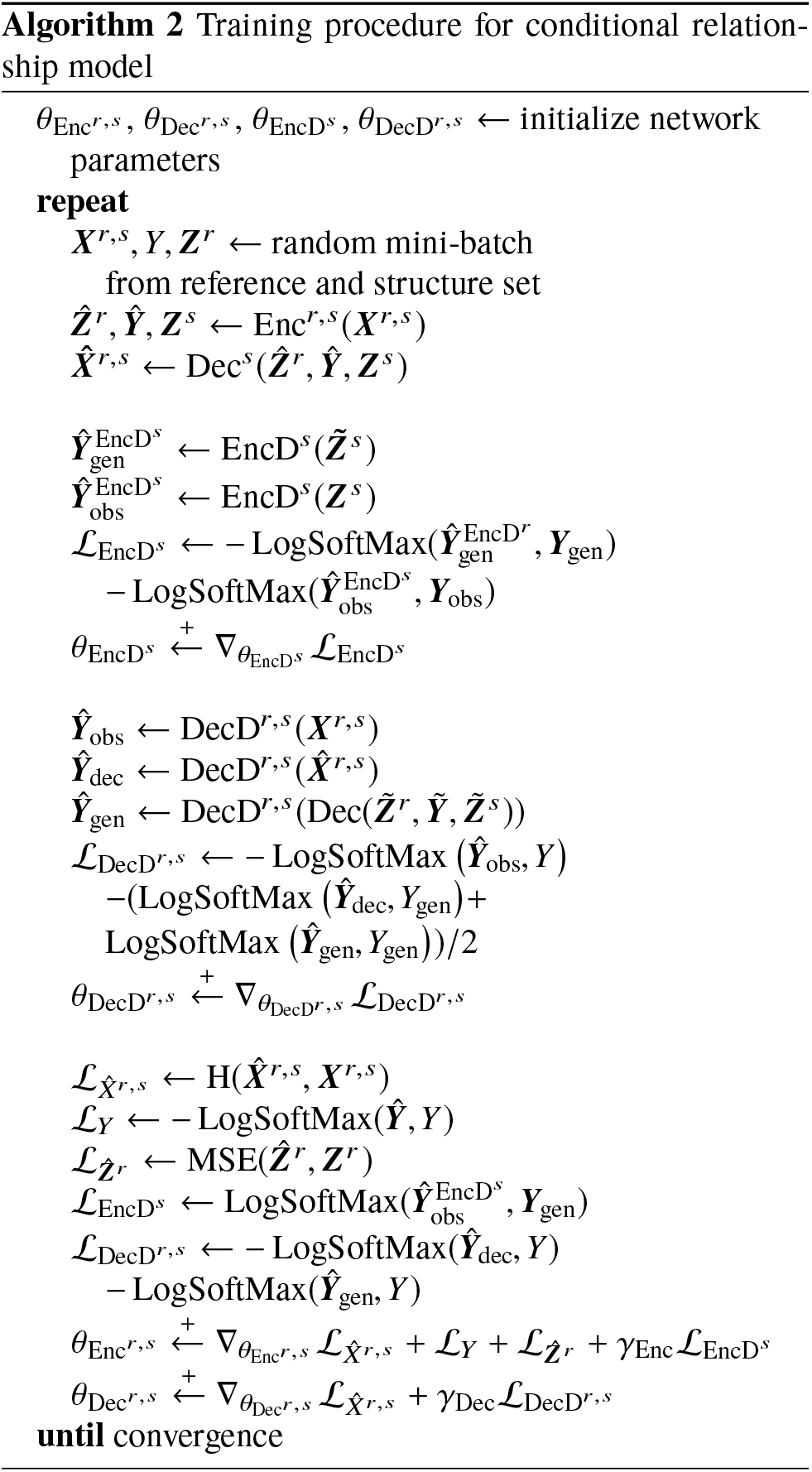

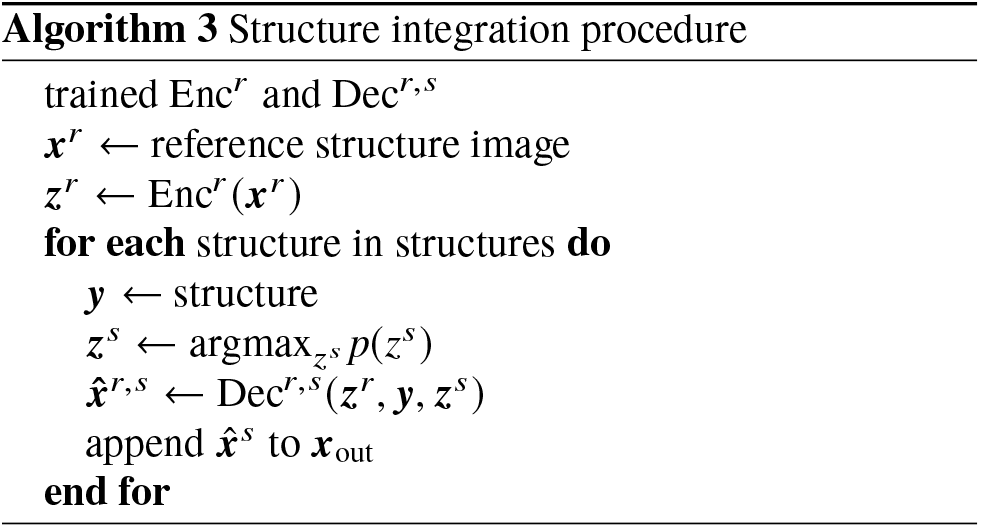

### 2.5. Estimate of structure variation

Given the differential relationships between these structures of interest and cell and nuclear shape, we are able to estimate the strength of the relationship between any given structure localization and the corresponding cell and nuclear shape. That is, if we fix the cell and nuclear shape embedding, *z^r^*, and structure type, ***y***, we are able to measure how much variability we observe as we sample from the structure localization embedding distribution, *z^s^*, resulting an an estimate of structure variation.

## 3. Results

### 3.1. Data Set

We use a collection of 3D segmented cell images generated from static 3D spinning disc confocal microscopy images of human induced pluripotent stem cells. These cells are from clonal lines, each gene edited to express mEGFP on a protein that localizes to a specific structure of interest, e.g. α-actinin (actin bundles), α-tubulin (microtubules), β-actin (actin filaments), desmoplakin (desmosomes), fibrillarin (nucleolus), lamin B1 (nuclear membrane), myosin IIB (actomyosin bundles), ST6GAL1 (golgi), Sec61β(endoplasmic reticulum), TOM20 (mitochondria), and ZO1 (tight junctions). Details on the cell lines, microscopy pipeline, and on the source image collection are available via the Allen Cell Explorer at http://www.allencell.org. Briefly, each image consists of channels corresponding to the nuclear signal, cell membrane signal, and a labeled sub-cellular structure of interest (Figure 2). Individual cells were segmented from a field, and each channel was processed by subtracting the most populous pixel intensity, zeroing-out negative-valued pixels, and rescaling image intensity to a value between 0 and 1. The cells were aligned by the major axis of the cell shape, centered according to the center of mass of the segmented nuclear region, and flipped according to image skew. Each of the 21,967 cell images were rescaled to 0.317 μm/px, and padded to 128 × 96 × 64 cubic voxels.

### 3.2. Model implementation

A summary of the model architectures is offered in Section C. The explicit model definitions and training procedures are available in the source code at https://github.com/AllenCellModeling/pytorch_integrated_cell.

We based the architectures and their implementations on a combination of resources, primarily (Larsen et al.,2015; Makhzani et al., 2015; Radford et al., 2015), and Kai Arulkumaran’s Autoencoders package (Arulkumaran, 2017).

We found that adding white noise to the first layer of decoder adversaries, DecD*^r^* and DecD*^r,s^*, stabilizes the relationship between the adversary and the autoencoder and improves convergence as in (Sønderby et al., 2016) and (Salimans et al., 2016).

We choose a 128-dimensional latent space for both ***Z**^r^* and ***Z**^s^*.

### 3.3. Training

To train the model, we used the Adam optimizer(Kingma & Ba, 2014) to perform gradient-descent, with a batch size of 30 and learning rate of 0.0002 for Enc*^r^*, Dec*^r^*, DecD*^r^*, Enc*^r,s^*, Dec*^r,s^*, DecD*^r,s^*, a learning rate of 0.01 for EncD*^r^* and EncD_s_, with *γ*_Enc_ and *γ*_Dec_ values of 10^−4^ and 10^−5^ respectively. The dimensionality of the latent spaces ***Z**^r^* and ***Z**^s^* were set to 128, and the prior distribution for both is an isotropic gaussian.

We split the data set into 95% training and 5% test (for more details see Table S9), and trained the model of cell and nuclear shape for 150 epochs, and the conditional model for 150 epochs.

The training curves for the reference and conditional model are shown in Figure S3.

The model was implemented in PyTorch (http://pytorch.org) version 0.1.12., and run on 2 NVIDIA V100 graphics cards via NVIDIA-docker. The model took approximately two weeks to train. Further details of our implementation can be found in the software repository (Section 4).

### 3.4. Experiments

We performed a variety of experiments to explore the utility of the model. While quantitative assessment is paramount, the nature of the data makes qualitative assessment indispensable, so we include assessments of this type in addition to more traditional measures of performance.

#### 3.4.1. Image Classification

While classification is not our primary use-case, it is a worthwhile benchmark of a well-functioning multi-class generative model. To evaluate the performance of the class-label identification of Enc*^r,s^*, we compared the results of the predicted labels and true labels on our hold out set. A summary of the results of our multinomial classification task is shown in Table S11. Our model is able to accurately classify most structures, and has trouble only on the poorly sampled or underrepresented classes.

#### 3.4.2. Image reconstruction

A necessary but not sufficient condition for the model to be useful is that cell images reconstructed from their latent space representations resemble the native images. Examples of image reconstruction from the training and test set are shown in Figure S1 for reference structures and Figure S2 for the structure localization model. The model recapitulates the essential structure localization patterns in the cells, and produce convincing reconstructions of cell and nuclear shape in both the training and test data.

#### 3.4.3. Integrating Cell Images

Conditional upon the cell and nuclear shape, the model predicts the most probable position of any particular structure via Algorithm 3. Some examples of integrating structure localization given cell and nuclear shapes is shown in Figure 3.

**Figure 3:**
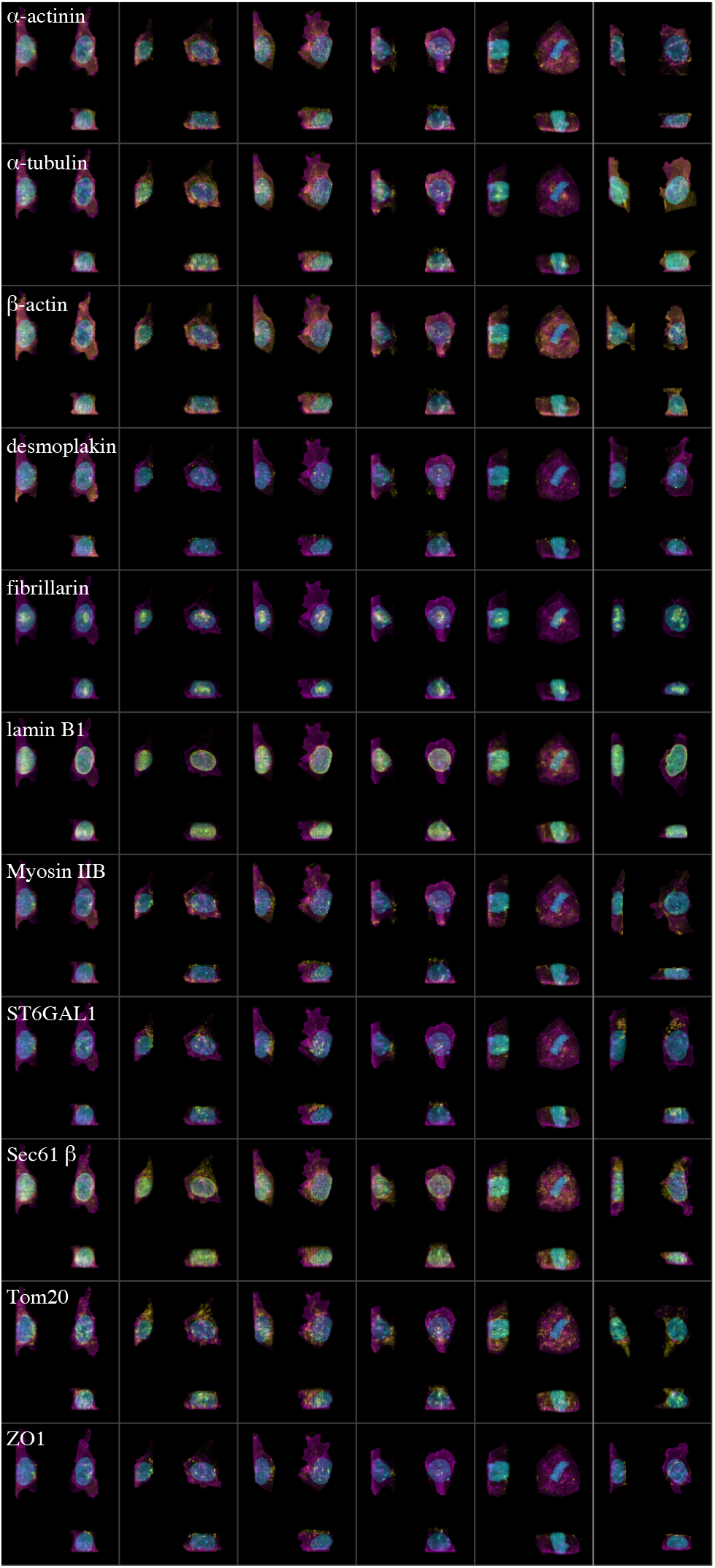
Most probable localization patterns predicted for selected cells for each structure (rows, top to bottom, structure as labeled, shown in yellow). The first five columns show the most probable localization for each structure, given the cell and nuclear shape. The last column (far right) shows an experimentally observed cell with that labeled structure for comparison. Reference structures, cell membrane and nucleus (DNA), are in magenta and cyan, respectively. Images have been cropped for visualization purposes. Note, for example, how fibrillarin resides within the DNA, and lamin B1 surrounds the DNA. See Figure S5b for structure channel only.

#### 3.4.4. Evaluation of Generated Image Distribution

To determine how well our model captures the variation in our data, we compare metrics computed on our input images and compare these to metrics computed on model-generated images.

We choose two methods for evaluating this variation: a pixel-wise measure of image similarity and the so-called inception score (Salimans et al., 2016).

As our measure of image similarity, we compute the pixel-wise cross correlation between the structure channels, ***x***^s^**, of every image in our training set. The distribution of this correlation is reported in Figure S6a. As expected, we can see that there is a large difference in this correlation across structures. Stereotypically localized structure labels, such as lamin B1 and fibrillarin, correlate highly; punctate cytoplasmic labels, such as desmoplakin, ST5GAL1 and ZO1 are highly variable in their localization in the current model.

We compare these distributions to the pair-wise cross correlation of encoded-decoded images from both the train and test set (Figures. S6c, S6d), as well as images generated by our model de novo (Figure S6b). Comparing the data distribution to the encoded-decoded images in the training set (Figure S6c), we see that the model does not capture the total amount of variation of the data. This is expected, as the autoencoder can be interpreted as a lossy compression method. While these distributions differ slightly, direct inspection of these images can be seen in structure channel of Figure S2, and these appear to be qualitatively similar.

Comparing the correlation between sampled images and the train and test images in (Figures S6c, S6d, S6b), we see that the distributions of the generated images are very similar to the encoded-decoded held-out test data, indicating that the model accurately captures high-level variation in the data in a generalizable manner.

An alternate evaluation metric is via the “inception score” introduced above (Salimans et al., 2016):

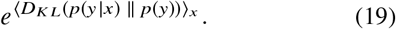

The inception score is the exponential of the average over the input data of the *KL*-divergence of the conditional probability of a label given an image, with respect to the marginal probabilities of each label. It a widely used measure of performance for generative models.

To compute the inception scores, we pass the generated images through the encoder Enc*^r,s^* and record the predictions for the class of each image, *p*(*y* | *x*), from which the marginal *p*(*y*) is trivial. If the generator Dec*^r,s^* creates representative images, the estimated class of each image image *p*(*y* | *x*) should have low entropy, while the distribution of class probability across all images, *p*(*y*), should have high entropy.

In Table S10, we report inception scores for our train and test data, and the generated images for each class sampled at the train and test set frequencies. On the test set, our model achieved an inception score of 7.561, compared to a score of 7.823 on the encoded-decoded test data, and a score of 7.876 for raw test data itself.

While the absolute magnitude of these scores is not directly evaluable at present, the similarity of these scores indicates that the variety and quality of the generated structures is on par with those in the data distribution.

#### 3.4.5. Variation in Structure Localization

Given that the model reasonably approximates the data distribution, as described in the previous section, we are interested in how predictive the cell and nuclear shape is of structure localization. That is, how much intrinsic variation is there in the localization of a given structure, and how much of that structure’s localization is dictated by the shape of the cell and nucleus? This relationship is quantified to determine the model’s performance in predicting the location and variation of different subcellular structures in the following section.

We evaluated the variation the model captures by generating 1000 structure images per cell and nuclear shape, and assessing the variation in that structure localization (see4) by computing the negative log-determinant of the 1000x1000 matrix of pairwise correlations. By exploring this distribution of structure localization, we assessed the conditions in which structure organization is tightly coupled to that cell and nuclear shape. That is, for a given cell shape, the localization of one structure type may be highly variable as one moves through the structure latent space, while the location of another may not be as variable. Furthermore, it can be imagined that these relationships would vary from cell state to cell state.

We report the distribution of per-cell summary statistics for cell and nuclear shapes in our train and test set in Figure S7.

In addition to assessing structure-wise variability and accuracy, we can also investigate the relationships between these metrics and the cell cycle. Using expert annotation for a subset of 7592 of the cells, which annotate each cell as in one of eight phases of the cell cycle ({interphase, prophase, prophase 2, pro metaphase 1, pro metaphase 2, metaphase, anaphase, telophase-cytokinesis}), we subdivide our population in cells in stages, those that are labeled in as interphase, and those that are not. As seen in Figure S7, the distributions of variation per structure show interesting biological correlation: in mitosis, we see that there is more variation (shift toward smaller negative log-determinant) in the localization of lamin B1, fibrillarin, ST6GAL1, and Tom20, while the distribution of α-tubulin is less variable. From a biological point of view, this makes sense: the nuclear envelope breaks down in mitosis, entailing less stereotypy in the location of lamin B1 and fibrillarin, with corresponding fragmentation of mitochondria (Tom20) and Golgi (ST6GAL1), while α-tubulin coalesces in to spindles that are highly stereotyped in location.

On a per-structure basis, we are also able to sort the cells in our input data according to which cell and nuclear shape corresponds to both the most and the least varied structure predictions, as shown in Figure S8. Structures such as lamin B1, fibrillarin, ST6GAL1, and Tom20 show a clear enrichment for mitotic cells in the most variable structure predictions, as compared to the least variable predictions, while the opposite is true of α-tubulin. This example for mitosis suggests that the present model is able to learn how the variability in the localization of different subcellular structures may be coupled to cell state.

#### 3.4.6. Latent space traversal

We further explore the generative capacity of our model by mapping out the variation in cell morphology and structure localization via a traversal of the reference latent space with respect to cells in ordered stages of mitosis as introduced in the previous section.

Given the expert manual annotation of cell cycle for 7592 of our images and their corresponding cell and nuclear shape representations, *z^r^*, we select medoid landmark cells chosen from the test data as representative of each ordered stage in mitosis.

In our traversal of the 128 dimensional latent space of cell and nuclear shape, we interpolate linearly in radial coordinates between each medoid of our annotated stages, from interphase through metaphase. Samples along this path are shown in Figure S9.

As we move between stages of the cell cycle in the reference latent space, we also predict the localization of each of our structures of interest by generating the most probable structure channel localizations from the center of the structure latent space.

Although these cell cycle expert annotations were not used during model training, they offer an opportunity to investigate how well the model generalizes to unseen data along biologically important axes of variation. These predictions of localization demonstrate smooth transitions and plausible localization patterns as the cell shape progresses from interphase through metaphase on held-out data, demonstrating the ability of our model to capture and predict complex relationships in structure organization and potentially other key biological phenomena given new data.

## 4. Discussion

Building models that capture relationships between the morphology and organization of cell structures is difficult but important challenge. While previous research has focused on constructing subcellular structure-specific parametric approaches, due to the extreme variation in localization among different subcellular structures, these approaches may not be convenient to employ for all structures under all conditions. Here, we have presented a nonparametric conditional model of structure organization that generalizes well to a wide variety of localization patterns, encodes the variation in cell structure and organization, allows for a probabilistic interpretation of the image distribution, and generates high quality three-dimensional, integrated synthetic images.

Our model of cell and subcellular structure differs from previous generative models (Zhao & Murphy, 2007; Peng & Murphy, 2011; Johnson et al., 2015): we directly model fluorescent label localization, rather than the detected objects and their boundaries. While object segmentation can be essential in certain contexts, and helpful in others, when these approaches are not necessary, it can be advantageous to omit these non-trivial intermediate steps. Our model does not constitute a “cytometric” approach (i.e. counting objects), but since we are directly modeling the localization of signal, we drastically reduce the modeling time by minimizing the amount of segmentation and the task of evaluating this segmentation with respect to the “ground truth”.

Even considering these differences, the model is compatible with existing frameworks and allows for mixed parametric and non-parametric localization relationships, since the model can be used for predicting the localization of structures when an appropriate parametric representation may not exist.

Since the post of our initial preprint manuscript and software (Johnson et al., 2017), there has been increased interest in the application of GANs to modeling cell morphology and organization. (Osokin et al., 2017) constructed a network that can generate images of cell shape, and generates additional subcellular structures conditioned on those. (Goldsborough et al., 2017) uses a GAN to generate images of cells across different conditions and evaluate the performance of a discriminator in identifying the mechanism of action of these conditions. These other GAN approaches (Osokin et al., 2017; Goldsborough et al., 2017), allow neither a probabilistic interpretation nor prediction of structure localization, in *real* images. Furthermore, the methods presented above generate 2D images with one to two orders of magnitude more training data than the method presented here. Finally, in contrast to these models, our method allows for a reversible mapping of images to a low-dimensional latent space, which permits the prediction of novel subcellular localization patterns in extant data.

Our model permits several straightforward extensions, including extension to time series. Because of the flexibility of our latent-space representation, we can potentially explicitly encode information regarding known cell states and/or perturbations, such as position in cell cycle, time after drug treatment, or “distance” along a differentiation pathway. Given sufficient information, it would be possible to encode a representation of a “structure space” to predict the localization of unobserved structures, or “perturbation space”, as in (Paolini et al., 2006). Coupling this with active learning approaches (Naik et al., 2016) opens the way to potentially build models that learn and encode the localization of diverse subcellular structures under a variety different conditions.

### Software and Data

The code for running the models used in this work is available at https://github.com/AllenCellModeling/pytorch_integrated_cell

The data used to train the model is available at s3://aics.integrated.cell.arxiv.paper.data.

## Acknowledgements

We would like to thank Robert F. Murphy, Julie Theriot, Rick Horwitz, Graham Johnson, Forrest Collman, Sharmishtaa Seshamani and Fuhui Long for their helpful comments, suggestions, and support in the preparation of the manuscript.

Furthermore, we would like to thank all members of the Allen Institute for Cell Science team, who generated and characterized the gene-edited cell lines, developed image-based assays, and recorded the high replicate data sets suitable for modeling. We particularly thank Liya Ding for segmentation data and Irina Mueller for mitosis annotations. These contributions were absolutely critical for model development.

We would like to thank Paul G. Allen, founder of the Allen Institute for Cell Science, for his vision, encouragement and support.

## Author Contributions

GRJ and RMD conceived, designed and implemented experiments. GRJ, RMD, and MMM wrote the paper.

## A. Supplementary Figures

**Figure S1:**
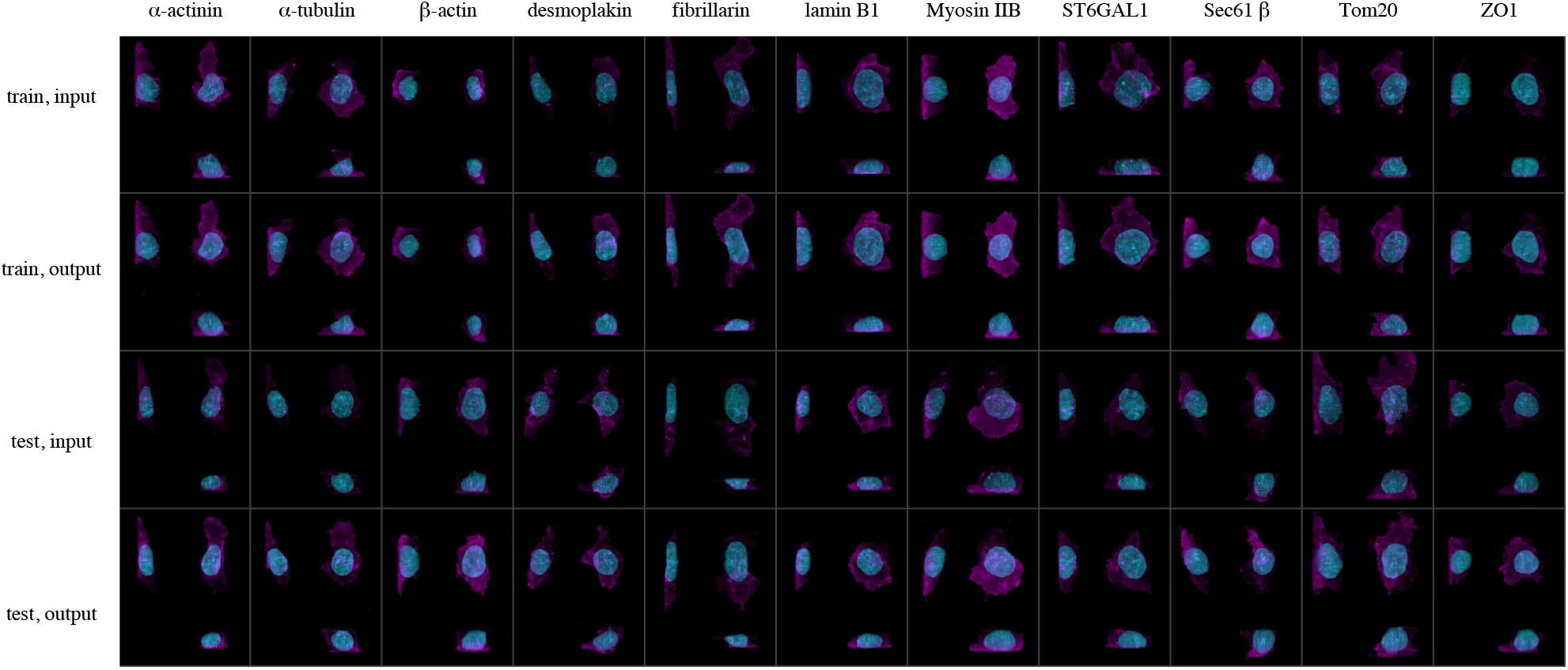
Image input (rows 1 and 3) and reconstruction (rows 2 and 4) from the reference model, showing training set (above two rows), and test set (bottom two rows).

**Figure S2:**
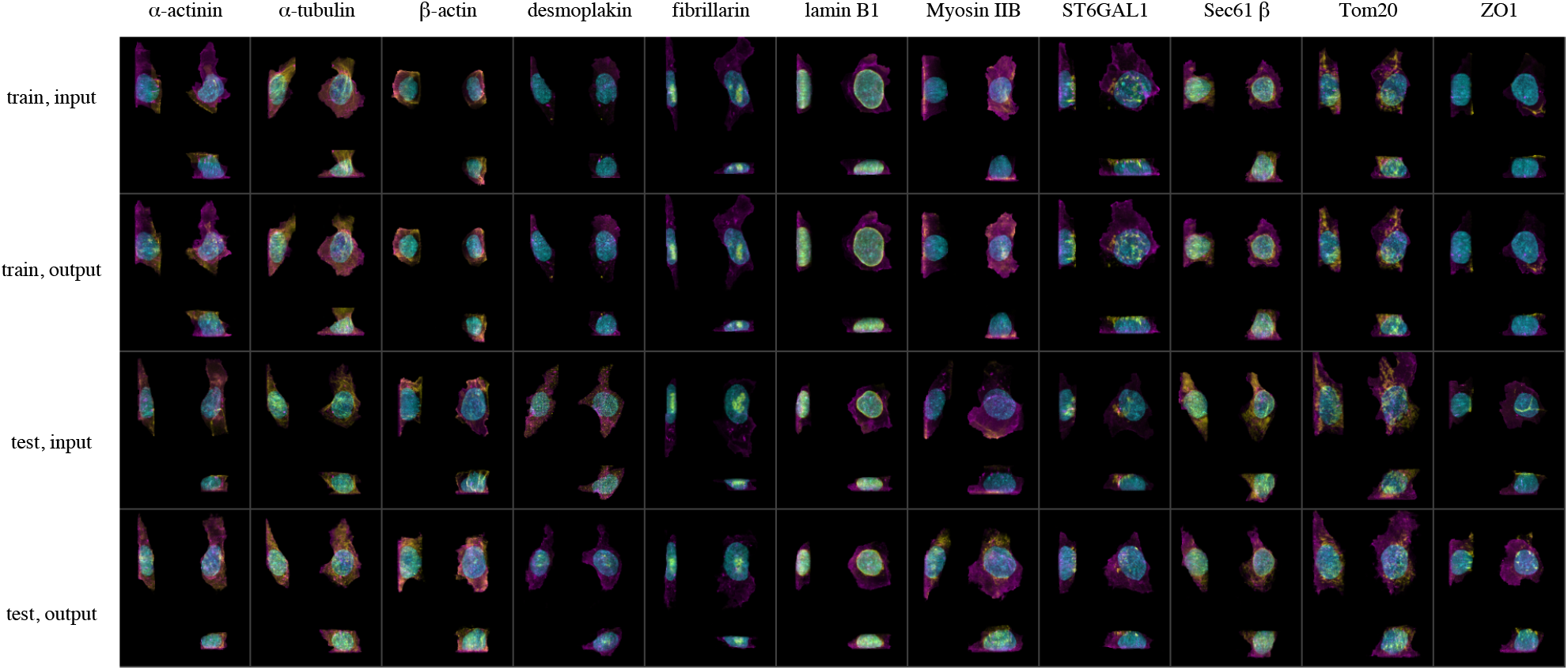
Image input (rows 1 and 3) and reconstruction (rows 2 and 4) from the structure model, showing training set (above two rows), and test set (bottom two rows).

**Figure S3:**
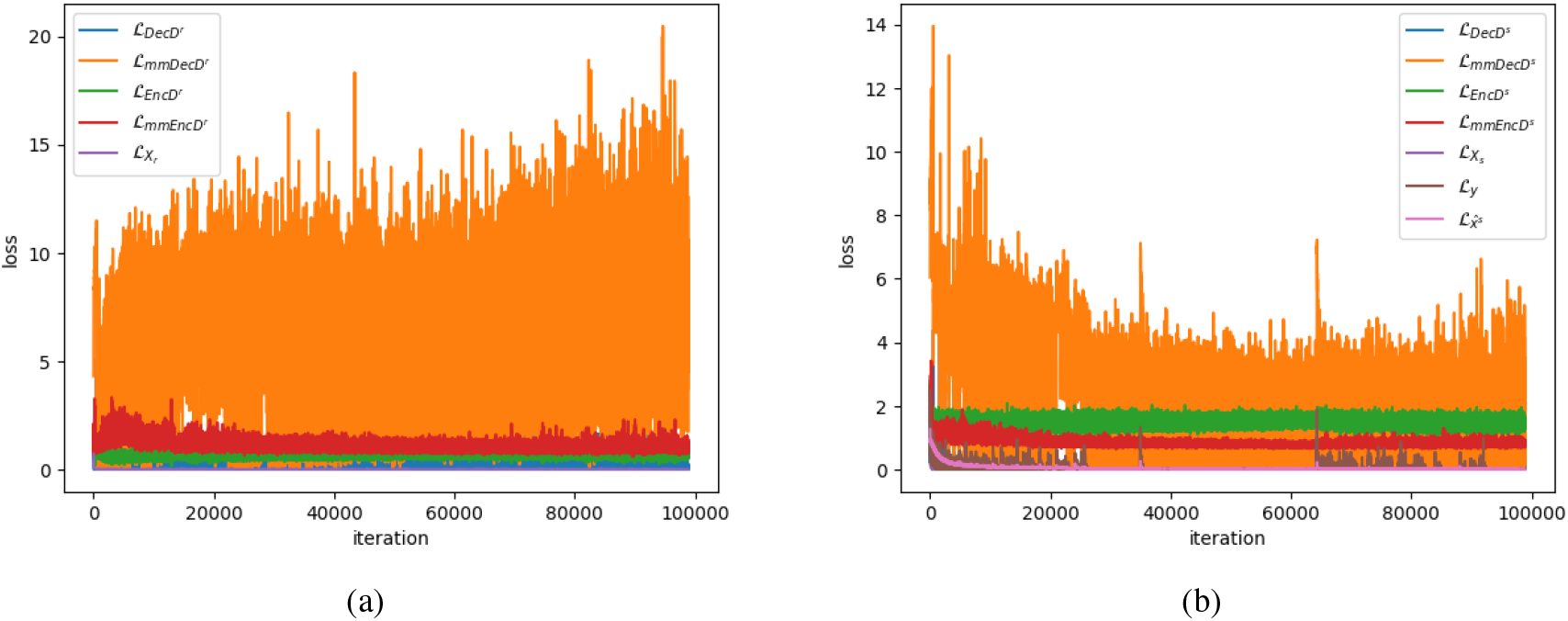
Training curves for the training of the reference model (a) and conditional model (b)

**Figure S4:**
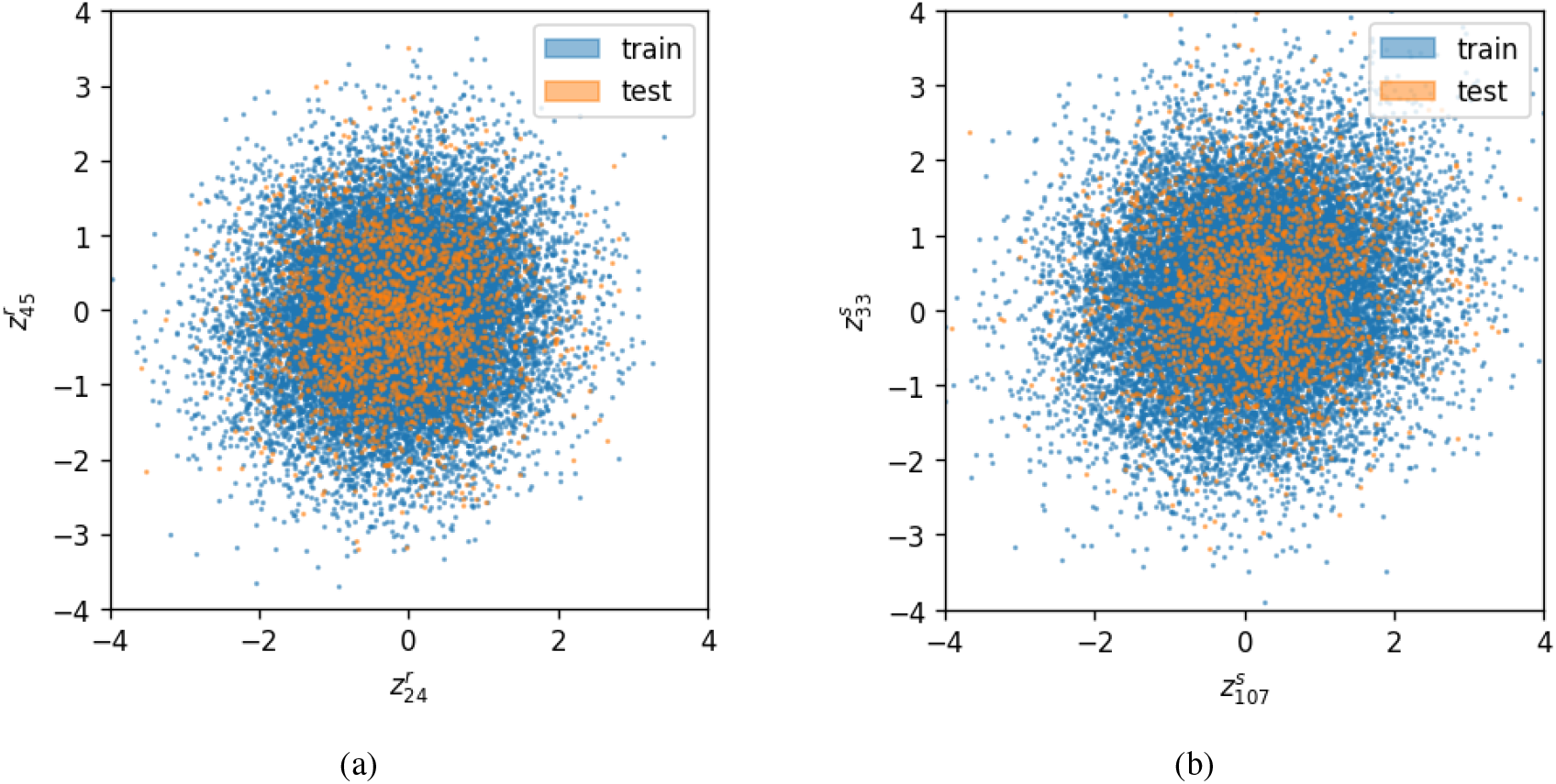
(a) two randomly selected dimensions of the reference structure latent space ***Z**^r^*. (b) two randomly selected dimensions of the reference structure latent space ***Z**^s^*.

**Figure S5:**
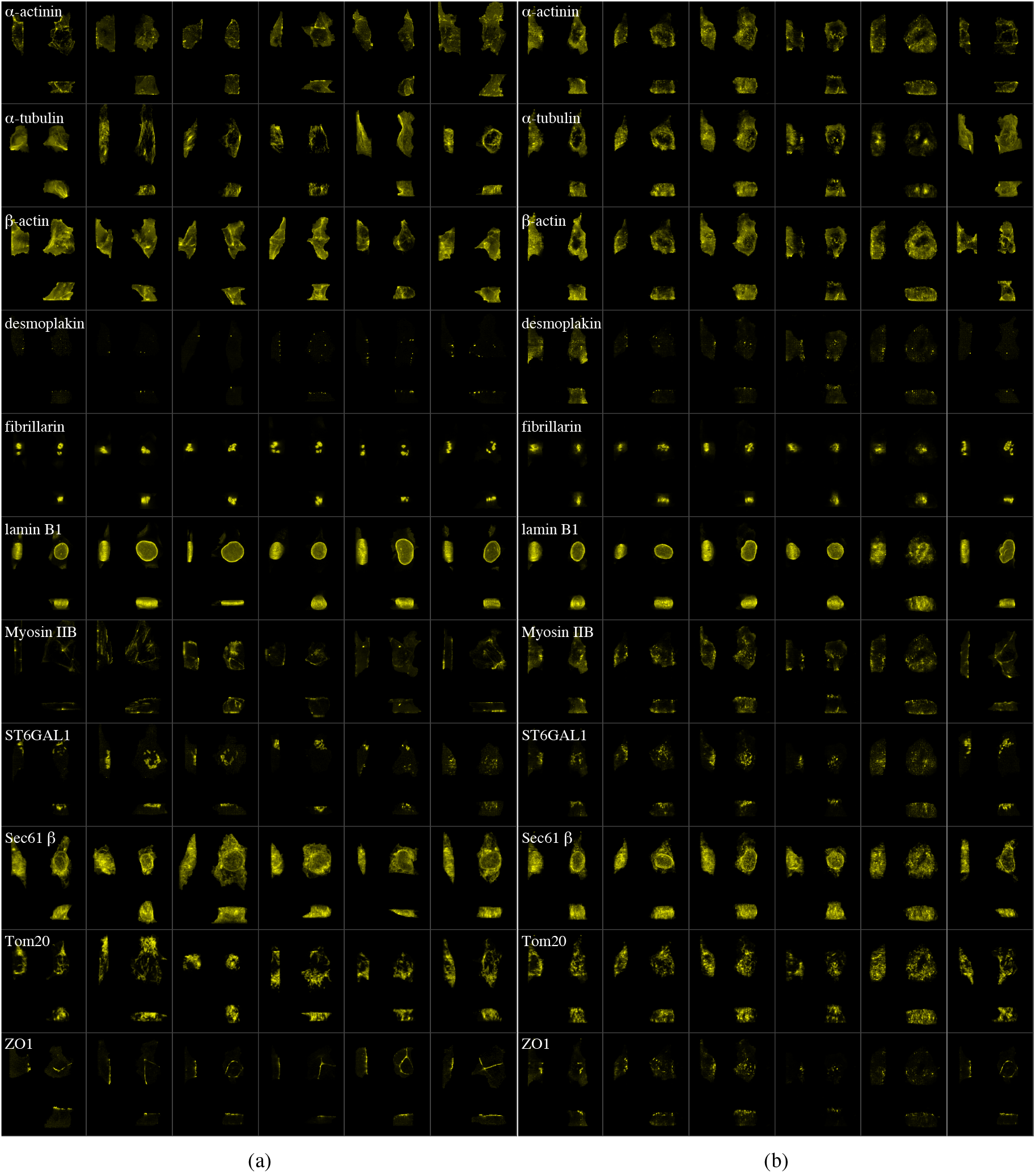
(a) Example structure channels for each of the 10 labeled structures in this paper and (b) predicted most probable localization patterns for selected cells from each labeled pattern. The first 5 columns show the most probable localization for the corresponding structures given the the same cell and nuclear shape. The last column shows a observed cell with that labeled structure. Rows correspond to structure types. Images have been cropped for visualization purposes.

**Figure S6:**
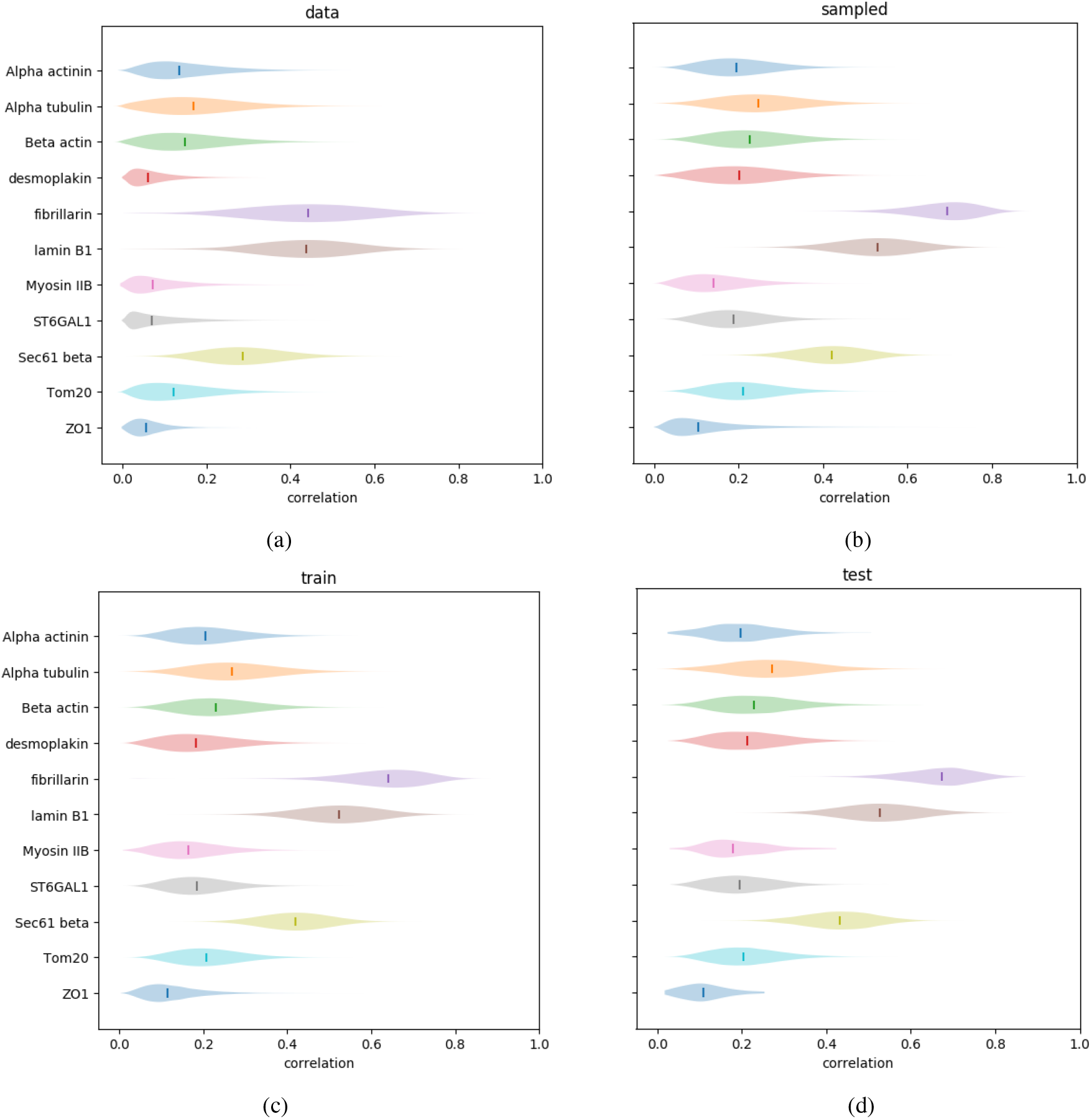
Distribution of pixel-wise correlation between all pairs of structure channels across image collections. (a) complete set of image data. (b) 2000 images sampled from the latent space distributions for each structure. (c) All encoded-decoded images in the training set. (d) All encoded-decoded images in the test set.

**Figure S7:**
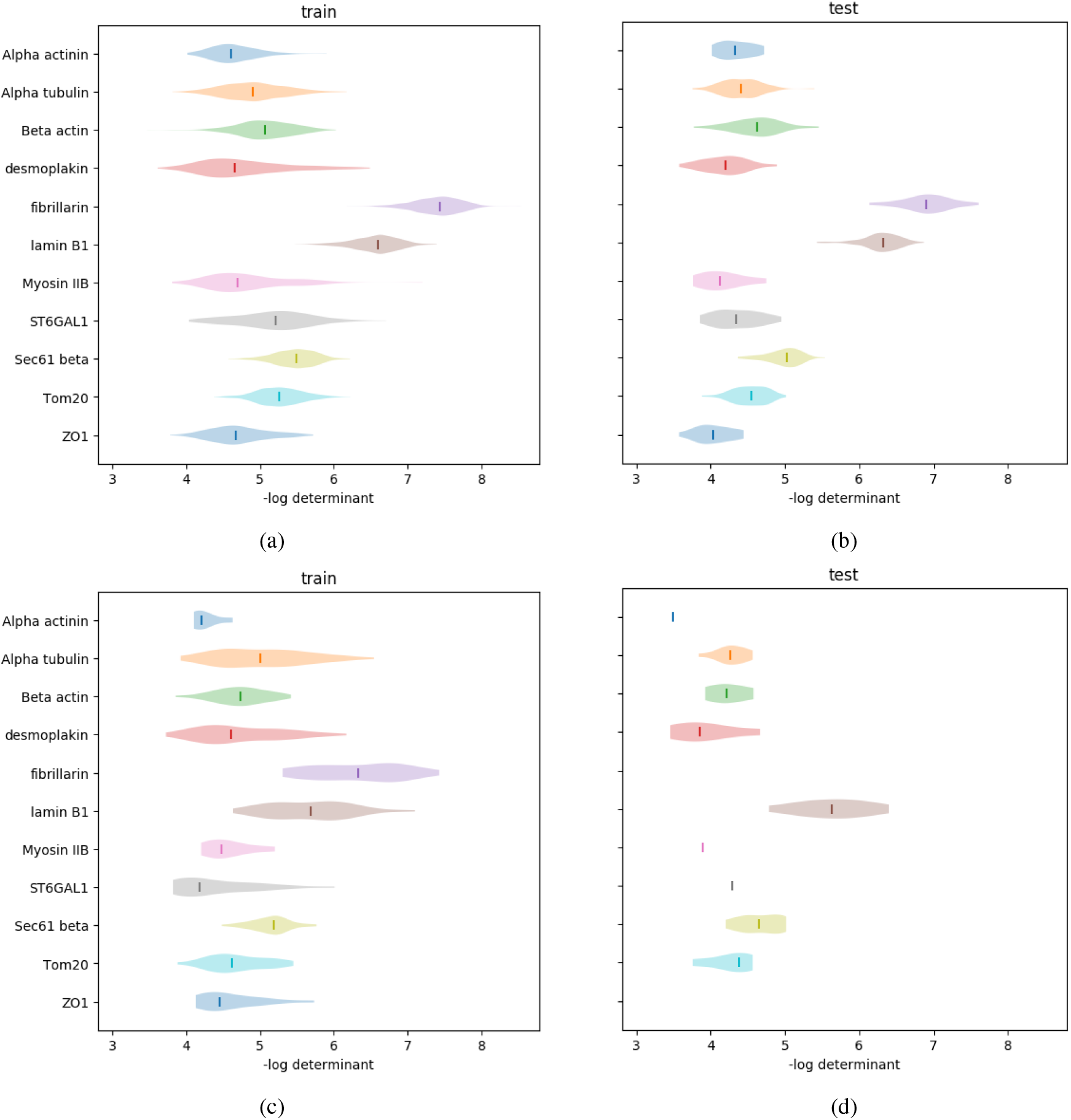
Distribution of per-cell variation statistics. For each cell and nuclear shape, we compute the variation between all pairs of 1000 generated structure channels in a 1000-by-1000 matrix, and compute the log determinant of that matrix. (a) training images labeled as interphase. (b) testing images labeled as interphase. (c) training images labeled as any stage of mitosis. (d) test images labeled as any stage of mitosis.

**Figure S8:**
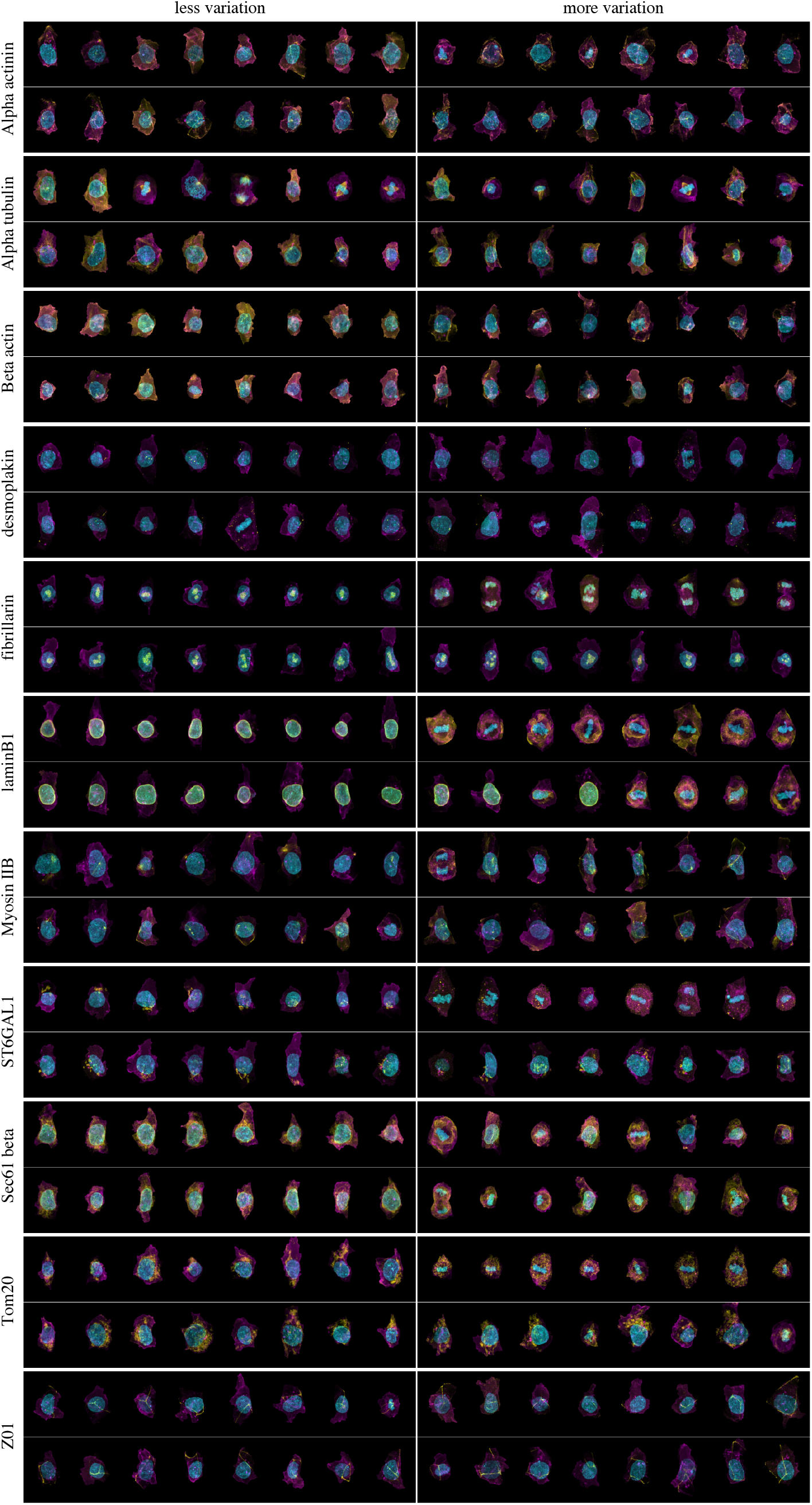
For each structure, we show the cells in our input data (top:train, bottom:test) that are predicted to have the most (right) and least (left) amount of variation in structure localization, based upon morphology of the cell. We see a clear enrichment for mitosis in several of the structures, indicating that the model has achieved an ability to differentially couple the variability in localization of subcellular components across structures.

**Figure S9:**
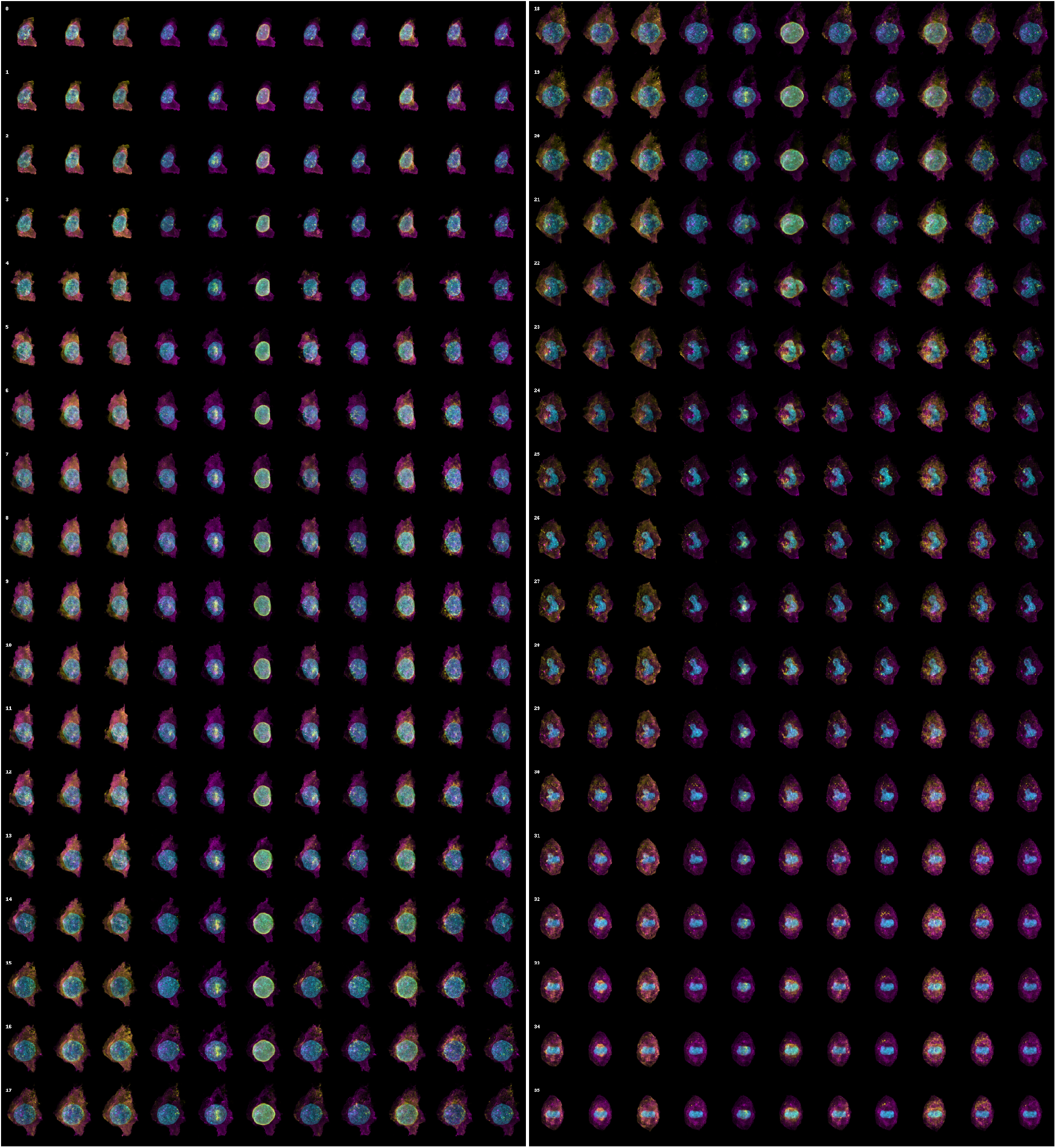
Generated cell and nuclear shapes and corresponding most probable predicted localization patterns for a walk from one mitotic state to the next. Each row indicates a time point, with columns corresponding to predicted localization patterns for each

## B. Supplementary Algorithms

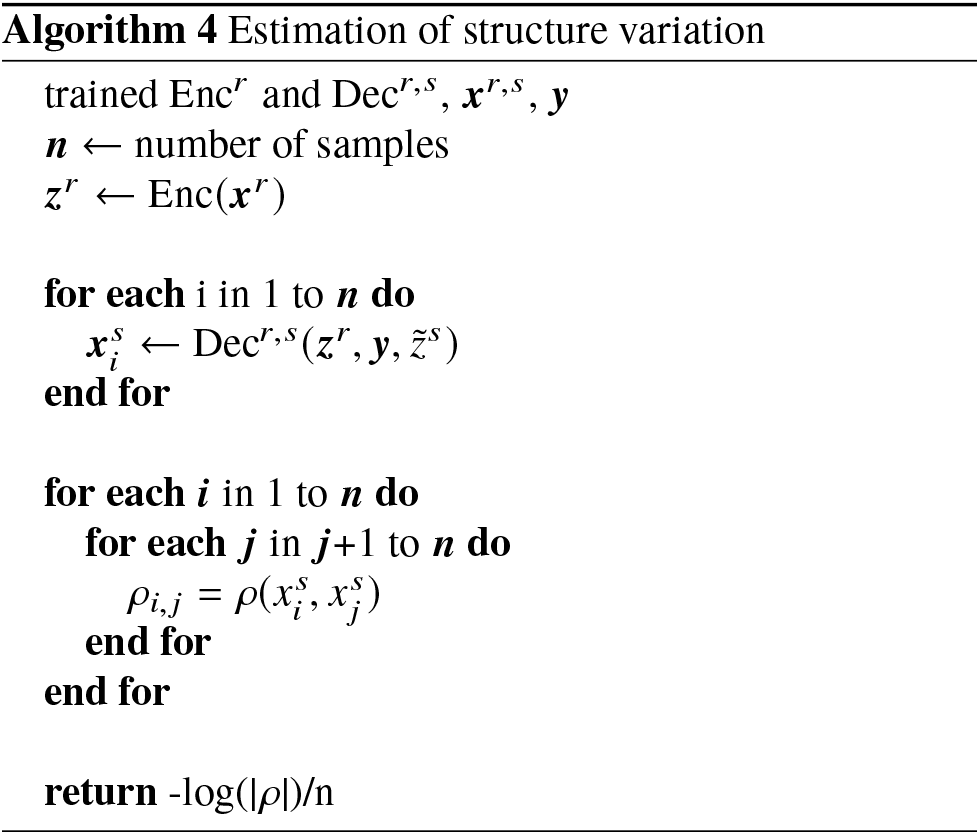

## C. Model Architectures

**Table S1:**
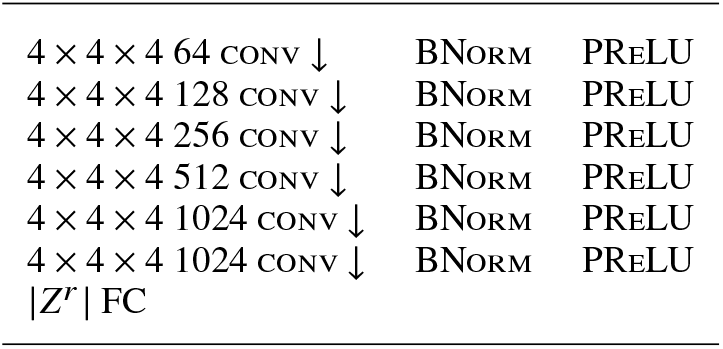
Architecture of Enc*^r^*

**Table S2:** Architecture of Dec*^r^*

|  |  |  |
| --- | --- | --- |
| 1024 FC | BNORM | PRELU |
| $4 \times 4 \times 4$ 1024 CONV ↑ | BNORM | PRELU |
| $4 \times 4 \times 4$ 512 CONV ↑ | BNORM | PRELU |
| $4 \times 4 \times 4$ 256 CONV ↑ | BNORM | PRELU |
| $4 \times 4 \times 4$ 128 CONV ↑ | BNORM | PRELU |
| $4 \times 4 \times 4$ 64 CONV ↑ | BNORM | PRELU |
| $4 \times 4 \times 4$ $\vert r\vert $ CONV ↑ | | SIGMOID |

**Table S3:** Architecture of EncD*^r^*

|  |  |  |
| --- | --- | --- |
| 1024 FC |  | LEAKY RELU |
| 1024 FC | BNORM | LEAKY RELU |
| 512 FC | BNORM | LEAKY RELU |
| 1 FC |  | SIGMOID |

**Table S4:** Architecture of DecD*^r^*

|  |  |  |
| --- | --- | --- |
| + WHITE NOISE $\sigma = 0.01$ | | |
| $4 \times 4 \times 4$ 64 CONV ↓ | | LEAKYRELU |
| $4 \times 4 \times 4$ 128 CONV ↓ | BNORM | LEAKYRELU |
| $4 \times 4 \times 4$ 256 CONV ↓ | BNORM | LEAKYRELU |
| $4 \times 4 \times 4$ 512 CONV ↓ | BNORM | LEAKYRELU |
| $4 \times 4 \times 4$ 512 CONV ↓ | BNORM | LEAKYRELU |
| $4 \times 4 \times 4$ 1 CONV ↓ | | SIGMOID |

**Table S5:**
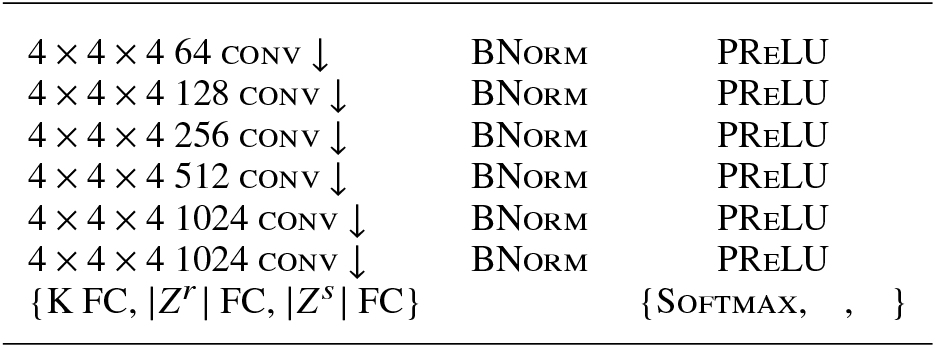
Architecture of Enc**^r,s^**

**Table S6:** Architecture of Dec*^r,s^*

|  |  |  |
| --- | --- | --- |
| 1024 FC | BNORM | PRELU |
| $4 \times 4 \times 4$ 1024 CONV ↑ | BNORM | PRELU |
| $4 \times 4 \times 4$ 512 CONV ↑ | BNORM | PRELU |
| $4 \times 4 \times 4$ 256 CONV ↑ | BNORM | PRELU |
| $4 \times 4 \times 4$ 128 CONV ↑ | BNORM | PRELU |
| $4 \times 4 \times 4$ 64 CONV ↑ | BNORM | PRELU |
| $4 \times 4 \times 4$ $\vert r + s\vert $ CONV ↑ | | SIGMOID |

**Table S7:** Architecture of EncD*^s^*

|  |  |  |
| --- | --- | --- |
| 1024 FC |  | LEAKY RELU |
| 1024 FC | BNORM | LEAKY RELU |
| 512 FC | BNORM | LEAKY RELU |
| K+1 FC |  | SIGMOID |

**Table S8:** Architecture of DecD**^r,s^**

|  |  |  |
| --- | --- | --- |
| + WHITE NOISE $\sigma = 0.01$ | | |
| $4 \times 4 \times 4$ 64 CONV ↓ | BNORM | LEAKYRELU |
| $4 \times 4 \times 4$ 128 CONV ↓ | BNORM | LEAKYRELU |
| $4 \times 4 \times 4$ 256 CONV ↓ | BNORM | LEAKYRELU |
| $4 \times 4 \times 4$ 512 CONV ↓ | BNORM | LEAKYRELU |
| $4 \times 4 \times 4$ 512 CONV ↓ | BNORM | LEAKYRELU |
| $4 \times 4 \times 4$ K+1 CONV ↓ | | SIGMOID |

## D. Data

**Table S9:**
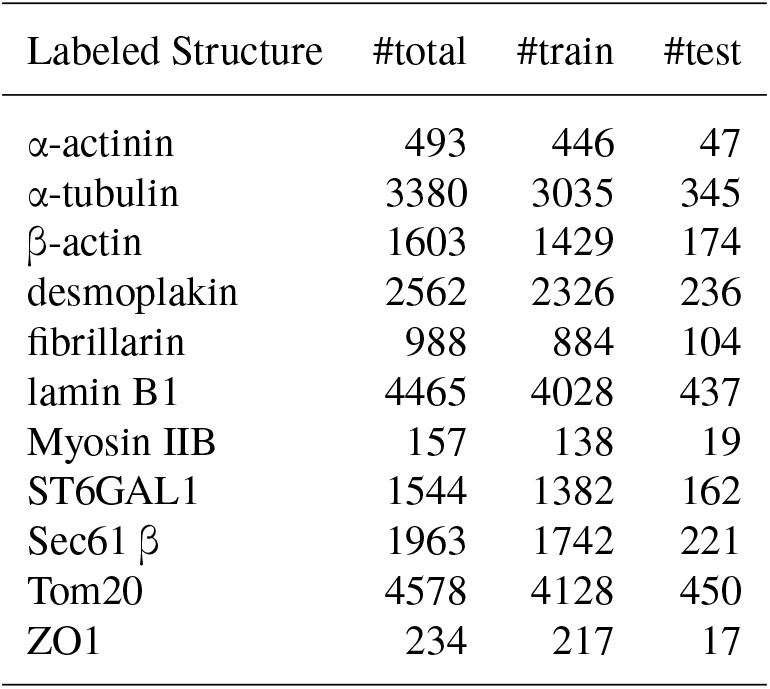
Labeled structures and their train/test split

| Labeled Structure | #total | #train | #test |
| --- | --- | --- | --- |
| α-actinin | 493 | 446 | 47 |
| α-tubulin | 3380 | 3035 | 345 |
| β-actin | 1603 | 1429 | 174 |
| desmoplakin | 2562 | 2326 | 236 |
| fibrillarin | 988 | 884 | 104 |
| lamin B1 | 4465 | 4028 | 437 |
| Myosin IIB | 157 | 138 | 19 |
| ST6GAL1 | 1544 | 1382 | 162 |
| Sec61 $\beta$ | 1963 | 1742 | 221 |
| Tom20 | 4578 | 4128 | 450 |
| ZO1 | 234 | 217 | 17 |

**Table S10:** Inception scores. KL divergence was computed with respect to *p*(*y*) across all classes.

| Labeled Structure | data |  | encoded-decoded |  | generated |  |
| --- | --- | --- | --- | --- | --- | --- |
|  | train | test | train | test | train | test |
| α-actinin | 44.29 | 34.904 | 44.252 | 36.209 | 28.316 | 32.082 |
| α-tubulin | 6.509 | 6.42 | 6.509 | 6.524 | 6.319 | 6.21 |
| β-actin | 13.824 | 11.929 | 13.822 | 12.434 | 12.47 | 11.684 |
| desmoplakin | 8.493 | 9.228 | 8.492 | 8.732 | 7.48 | 8.273 |
| fibrillarin | 22.347 | 20.904 | 22.347 | 20.899 | 22.07 | 21.168 |
| lamin B1 | 4.904 | 5.05 | 4.904 | 5.02 | 4.915 | 5.068 |
| Myosin IIB | 143.138 | 41.747 | 143.106 | 25.608 | 31.333 | 18.535 |
| ST6GAL1 | 14.294 | 13.106 | 14.294 | 12.867 | 13.223 | 12.808 |
| Sec61 $\beta$ | 11.34 | 9.687 | 11.34 | 9.843 | 11.103 | 9.884 |
| Tom20 | 4.786 | 4.869 | 4.786 | 4.892 | 4.788 | 4.877 |
| ZO1 | 91.031 | 83.076 | 91.008 | 54.921 | 13.355 | 11.081 |
| all classes | 8.085 | 7.876 | 8.085 | 7.823 | 7.493 | 7.561 |

**Table S11:** Predicted structure labels from Enc*^r,s^* on hold out

|  |  |  |  |  |  |  |  |  |  |  |  |
| --- | --- | --- | --- | --- | --- | --- | --- | --- | --- | --- | --- |
| α-actinin | 36 | 0 | 4 | 1 | 0 | 0 | 5 | 1 | 0 | 0 | 0 |
| α-tubulin | 1 | 330 | 5 | 1 | 2 | 2 | 1 | 0 | 2 | 1 | 0 |
| β-actin | 3 | 9 | 160 | 1 | 0 | 0 | 1 | 0 | 0 | 0 | 0 |
| desmoplakin | 0 | 0 | 0 | 229 | 0 | 0 | 0 | 4 | 0 | 0 | 3 |
| fibrillarin | 0 | 0 | 0 | 0 | 104 | 0 | 0 | 0 | 0 | 0 | 0 |
| lamin B1 | 0 | 0 | 0 | 0 | 0 | 434 | 0 | 1 | 0 | 2 | 0 |
| Myosin IIB | 4 | 0 | 0 | 5 | 0 | 0 | 7 | 0 | 0 | 0 | 3 |
| ST6GAL1 | 0 | 0 | 0 | 6 | 0 | 0 | 0 | 156 | 0 | 0 | 0 |
| Sec61β | 0 | 2 | 0 | 0 | 0 | 0 | 0 | 0 | 218 | 1 | 0 |
| Tom20 | 0 | 0 | 0 | 0 | 0 | 0 | 0 | 0 | 1 | 449 | 0 |
| ZO1 | 0 | 0 | 0 | 0 | 0 | 0 | 0 | 0 | 0 | 0 | 17 |

## References

Arulkumaran, Kai. Autoencoders, 2017. URL https://github.com/Kaixhin/Autoencoders.

Boland, Michael V and Murphy, Robert F. A neural network classifier capable of recognizing the patterns of all major subcellular structures in fluorescence microscope images of hela cells. Bioinformatics, 17(12):1213–1223, 2001.

Carpenter, Anne E, Jones, Thouis R, Lamprecht, Michael R, Clarke, Colin, Kang, In Han, Friman, Ola, Guertin, David A, Chang, Joo Han, Lindquist, Robert A, Moffat, Jason, Golland, Polina, and Sabatini, David M. CellProfiler: image analysis software for identifying and quantifying cell phenotypes. Genome biology, 7(10):R100, 2006.

Donovan, Rory M, Tapia, Jose-Juan, Sullivan, Devin P, Faeder, James R, Murphy, Robert F, Dittrich, Markus, and Zuckerman, Daniel M. Unbiased rare event sampling in spatial stochastic systems biology models using a weighted ensemble of trajectories. PLoS computational biology, 12(2):e1004611, 2016.

Goldsborough, Peter, Pawlowski, Nick, Caicedo, Juan C, Singh, Shantanu, and Carpenter, Anne. Cytogan: Generative modeling of cell images. bioRxiv, pp. 227–646, 2017.

Goodfellow, Ian J, Pouget-Abadie, Jean, Mirza, Mehdi, Xu, Bing, Warde-Farley, David, Ozair, Sherjil, Courville, Aaron, and Bengio, Yoshua. Generative Adversarial Networks. arXiv.org, June 2014.

Johnson, G R, Buck, T E, Sullivan, D P, Rohde, G K, and Murphy, R F. Joint modeling of cell and nuclear shape variation. Molecular Biology of the Cell, 26(22):4046–4056, November 2015.

Johnson, Gregory R, Donovan-Maiye, Rory M, and Maleckar, Mary M. Generative modeling with conditional autoencoders: Building an integrated cell. arXiv preprint arXiv:1705.00092, 2017.

Kim, Min-Sik, Pinto, Sneha M, Getnet, Derese, Nirujogi, Raja Sekhar, Manda, Srikanth S, Chaerkady, Raghothama, Madugundu, Anil K, Kelkar, Dhanashree S, Isserlin, Ruth, Jain, Shobhit, et al. A draft map of the human proteome. Nature, 509(7502): 575–581, 2014.

Kingma, Diederik P and Ba, Jimmy. Adam: A Method for Stochastic Optimization. arXiv.org, December 2014.

Larsen, Anders Boesen Lindbo, Sønderby, Søren Kaae, Larochelle, Hugo, and Winther, Ole. Autoencoding beyond pixels using a learned similarity metric. arXiv.org, December 2015.

Makhzani, Alireza, Shlens, Jonathon, Jaitly, Navdeep, Goodfellow, Ian, and Frey, Brendan. Adversarial Autoencoders. arXiv.org, November 2015.

Murphy, R F. Location proteomics: a systems approach to subcellular location. Biochemical Society transactions, 33(Pt 3):535–538, June 2005.

Naik, Armaghan W, Kangas, Joshua D, Sullivan, Devin P, and Murphy, Robert F. Active machine learning-driven experimentation to determine compound effects on protein patterns. eLife, 5:e10047, February 2016.

Osokin, Anton, Chessel, Anatole, Salas, Rafael E Carazo, and Vaggi, Federico. Gans for biological image synthesis. arXiv preprint arXiv:1708.04692, 2017.

Paolini, Gaia V, Shapland, Richard H B, van Hoorn, Willem P, Mason, Jonathan S, and Hopkins, Andrew L. Global mapping of pharmacological space. Nature biotechnology, 24(7):805–815, July 2006.

Peng, Tao and Murphy, Robert F. Image-derived, threedimensional generative models of cellular organization. Cytometry Part A, 79A(5):383–391, April 2011.

Radford, Alec, Metz, Luke, and Chintala, Soumith. Unsupervised Representation Learning with Deep Convolutional Generative Adversarial Networks. arXiv.org, November 2015.

Rajaram, Satwik, Pavie, Benjamin, Wu, Lani F, and Altschuler, Steven J. PhenoRipper: software for rapidly profiling microscopy images. Nature Methods, 9(7):635–637, June 2012.

Salimans, Tim, Goodfellow, Ian, Zaremba, Wojciech, Cheung, Vicki, Radford, Alec, and Chen, Xi. Improved techniques for training gans. In Advances in Neural Information Processing Systems, pp. 2234–2242, 2016.

Sønderby, Casper Kaae, Caballero, Jose, Theis, Lucas, Shi, Wenzhe, and Huszár, Ferenc. Amortised MAP Inference for Image Super-resolution. arXiv.org, October 2016.

Zhao, Ting and Murphy, Robert F. Automated learning of generative models for subcellular location: Building blocks for systems biology. Cytometry Part A, 71A(12): 978–990, 2007.

